# Mesolimbic and mesocortical pathways differentially support fentanyl-context associations

**DOI:** 10.1101/2025.11.26.690827

**Authors:** Annalisa Montemarano, Hajra Sohail, Laura B. Murdaugh, Kyrsten Derr, Farrah A. Alkhaleel, Logan D. Fox, Samiksha Pandey, Megan E. Fox

## Abstract

**Rationale:** Synthetic opioids like fentanyl are contributing to unprecedented overdose rates, yet the neural circuitry underlying fentanyl-associated behaviors remains poorly understood. The ventral tegmental area (VTA) projects to both the nucleus accumbens (NAc) and prefrontal cortex (PFC), forming distinct pathways that are implicated in drug-cue associations, though their specific roles in fentanyl-context encoding are not well defined.

**Objectives:** This study aimed to determine how VTA-NAc and VTA-PFC circuits contribute to fentanyl-context associations, and to assess the role of downstream dopamine receptor signaling in fentanyl context-seeking.

**Methods:** Male and female mice underwent fentanyl conditioned place preference (CPP; 0.2 mg/kg). We locally inhibited dopamine D1 or D2 receptors in NAc or PFC during CPP expression. We used fiber photometry calcium imaging to measure activity in VTA-NAc and VTA-PFC projection neurons, and chemogenetic inhibition to suppress activity during CPP expression.

**Results:** Fentanyl CPP expression was attenuated by blocking D1 but not D2 receptors in PFC, and D2 but not D1 receptors in NAc. We found both VTA-NAc and VTA-PFC exhibited increased calcium activity during fentanyl exposure and during entries to the fentanyl-paired context. We further identified a functional role for VTA-NAc, as chemogenetic inhibition of VTA-NAc, but not VTA-PFC, reduced fentanyl context-seeking.

**Conclusions:** While both VTA-NAc and VTA-PFC pathways are recruited by fentanyl exposure, fentanyl context-seeking relies on different downstream dopamine receptors in NAc vs PFC. Further, activity in VTA-NAc functionally supports the expression of fentanyl CPP. Together, these findings indicate that VTA circuits differentially contribute to fentanyl context-seeking.

## INTRODUCTION

Opioid use disorder (OUD) is a chronic, relapsing condition that impacts millions of individuals. The rate of opioid overdose deaths in the U.S. has dramatically escalated in recent years, primarily due to increased prevalence of potent synthetic opioids like fentanyl (Spencer et al. 2024). Fentanyl possesses unique pharmacological properties compared to other opioids: its high lipid solubility allows for rapid passage through the blood brain barrier (Kelly et al. 2023), it exhibits unique binding interactions with the mu opioid receptor (Sutcliffe et al. 2022), and distinct receptor selectivity as compared to morphine (Eshleman et al. 2020). Fentanyl exposure is associated with poorer treatment outcomes for OUD (Socias et al. 2022; Kawasaki et al. 2026), necessitating novel advances in treatment for OUD cases that involve fentanyl. Despite the increased prevalence of fentanyl misuse, an understanding of the neuroadaptations that underly fentanyl misuse remains lacking.

Advancing OUD treatment requires improving our understanding of how opioid-cue associations are formed, as recognition of drug cues contributes to craving and eventual relapse (Vafaie and Kober 2022). One of the most common preclinical models of drug-cue associations is conditioned place preference (CPP). During CPP, subjects develop drug-context associations over several conditioning sessions, then will preferentially seek out the drug-paired context in the absence of drug. A significant mediator of these processes is the ventral tegmental area (VTA), a heterogenous midbrain structure important for drug reward (Koob and Volkow 2010; Doyle and Mazei-Robison 2021). VTA activity and dopamine release in downstream nucleus accumbens (NAc) and prefrontal cortex (PFC) are closely linked to CPP. VTA calcium activity increases during entries to a cocaine-paired context (Calipari et al. 2017), and VTA inactivation with baclofen and muscimol attenuates cocaine place preference (Shinohara et al. 2017). Extracellular dopamine increases in NAc during entries to a heroin-paired context (O’Neal et al. 2022), and in both NAc and PFC during the expression of cocaine CPP (Kawahara et al. 2021). Dopamine D1 and D2 receptors differentially regulate CPP depending on the conditioned drug. For example, amphetamine CPP is attenuated by either NAc D1 or D2 blockade (Hiroi and White 1991), while neither D1 nor D2 blockade influences nicotine (Spina et al. 2006) or alcohol CPP (Young et al. 2014). Morphine CPP is attenuated by D1 but not D2 blockade in NAc (Shippenberg et al. 1993), however these mechanisms may not extend to the synthetic opioid fentanyl. Indeed, fentanyl vs morphine differentially alter striatal dopamine metabolism (Finlay et al. 1988), necessitating additional investigation on dopaminergic signaling mechanisms of fentanyl-context associations.

While VTA dopamine contributes to CPP, not all VTA dopamine neurons are functionally identical. Rather, they exhibit several distinct properties depending on whether they project to NAc or PFC, including different baseline electrophysiological (Lammel et al. 2008; Chaudhury et al. 2013; Friedman et al. 2014; Juarez and Han 2016) and molecular (Sesack et al. 1998; Lewis et al. 2001; Lammel et al. 2008) signatures, as well as different responses to opioids (Margolis et al. 2006; Simmons et al. 2019; Morison et al. 2025). As such, VTA dopamine neurons that project to NAc or PFC may also exert differential regulation over opioid-associated behaviors, but this has not been fully characterized *in vivo*. Though VTA terminals in NAc exhibit increased activity in response to cocaine CPP (Calipari et al. 2017), it is unknown if this function extends to mesocortical neurons, or to opioid CPP. Furthermore, dopamine is not necessary for opioid-context associations, since dopamine deficient mice (Hnasko et al. 2005) and D1-receptor knockout mice (Ting-A-Kee et al. 2013) can form opioid place preferences. Non-dopamine neurons in VTA may be important for opioid-context associations. Indeed, the VTA contains multiple neuron subtypes capable of releasing glutamate, GABA, or a combination of these (Phillips et al. 2022; Fox et al. 2024; Fitzgerald and Day 2025), and recent work indicates many behavioral functions classically attributed to VTA dopamine neurons can also be mediated by glutamate (Root et al. 2014; Wang et al. 2015; Zell et al. 2020; McGovern et al. 2021), GABA (Tan et al. 2012; van Zessen et al. 2012), and co-transmitting (Yoo et al. 2016; Root et al. 2020; Warlow et al. 2024) neurons. These functions extend to opioid use, as VTA GABA and glutamate neurons express mu opioid receptors (Oriol et al. 2025), participate in opioid seeking behaviors (Bossert et al. 2004; McGovern et al. 2023), and undergo diverse molecular adaptations following fentanyl self-administration (Fox et al. 2024). An understanding of how multiple VTA neuronal subtypes contribute to opioid CPP can improve our understanding of how opioid-context associations are formed.

In this study we sought to understand how mesolimbic (VTA-NAc) and mesocortical (VTA-PFC) pathways differentially promote fentanyl context-seeking. Consistent with other drug-context associations, we found blocking D1 but not D2 receptors in PFC attenuated fentanyl CPP expression. Unlike other drug-context associations, we found blocking D2 but not D1 receptors in NAc attenuated fentanyl CPP expression. To investigate upstream VTA activity that could mediate these effects, we then targeted VTA neurons that either project to PFC or to NAc in a cell type-agnostic manner. Using *in vivo* calcium imaging, we found both projections exhibited similar activation during fentanyl CPP. However, chemogenetic inhibition of VTA-NAc, but not VTA-PFC, attenuated fentanyl context-seeking. These experiments support projection-specific functions for multiple VTA neuron subtypes in fentanyl context-seeking, thus providing mechanistic insight into the neural encoding of fentanyl-associated behaviors.

## MATERIALS AND METHODS

### Experimental subjects

All procedures were approved by the Institutional Animal Care and Use Committee at the Pennsylvania State University College of Medicine (PSUCOM) and conducted in accordance with NIH guidelines for the use of laboratory animals. C57BL/6J and RiboTag mice on a C57BL/6J background were obtained from The Jackson Laboratory (RRID:IMSR_JAX:029977) and bred at PSUCOM. All mice were given food and water *ad libitum* and housed in the PSUCOM vivarium on a 12:12h light:dark cycle with lights on at 7:00 am. Mice were group housed in corncob bedding and provided with nestlets, or pair-housed across a perforated acrylic divider after fiber optic or infusion cannula implantations to prevent isolation stress. All experiments used mice of both sexes (defined gonadally) aged 8-10 weeks at the start of behavior.

### Stereotaxic surgery

Adeno-associated viruses for Cre recombinase (AAVrg.hSyn.HI.eGFP-Cre.WPRE.SV40 or AAV-CMV-Pl-Cre), Cre-dependent GCaMP8s (AAV9-syn-FLEX-jGCaMP8s-WPRE), Cre-dependent Gi-coupled (AAV9-hSyn-DIO-hM4D(Gi)-mCherry) Designer Receptors Exclusively Activated by Designer Drugs (DREADDs), and Cre-dependent mCherry control (AAV9-hSyn-DIO-mCherry) were acquired from Addgene (viral preps #105540-AAVrg, 105537-AAV9, 162377-AAV9, 44362-AAV9, 50459-AAV9). Mice were anesthetized with 1-4% isoflurane gas in oxygen delivered at 1 liter/min, and affixed in a stereotaxic frame (Stoelting, Dale IL). Mice had a small burr hole made over target brain regions using the following coordinates (in mm, relative to bregma): NAc core (AP +1.6, ML ±1.5, DV -4.4, 10° angle), PFC (AP +1.7, ML ±0.75, DV -2.5, 15° angle), VTA (AP -3.2, ML ±1.0, DV -4.6, 7° angle). Viral vectors were delivered with Hamilton neurosyringes with 33-gauge needles at a rate of 100 nL/min for a total volume of 300 nL/virus, then needles were left in place for 5 min to minimize spread up the tract. For fiber photometry, AAV-CMV-Cre was infused bilaterally in NAc core or PFC of wildtype mice, 0.4 nL DIO-GCaMP was infused unilaterally in the right VTA (AP -3.2, ML +0.4, DV -4.2, 0° angle), two screws were fixed to the skull, then a fiber optic cannula (Thorlabs #CFM14L05) was implanted above the right VTA (DV -4.0) and fixed to the skull with C&B Metabond Adhesive Cement (Parkell #S380). For chemogenetic experiments, AAVrg-hSyn-eGFP-Cre was infused bilaterally in NAc core or PFC of wildtype mice, and DIO-hM4Di or DIO-mCherry was infused bilaterally in VTA. For Riboseq, AAVrg-hSyn-eGFP-Cre was infused bilaterally in NAc core or PFC of Ribotag mice. For intracranial pharmacology, two screws were fixed to the skull, then a bilateral guide cannula targeting PFC (RWD, Sugar Land, TX) was implanted at AP +2.1, ML ±0.3, DV -1.7, or targeting NAc (P1 Technologies, Roanoke, VA) was implanted at AP +1.6, ML ±1.5, DV -4.0. Cannula were fixed to the skull with dental cement (#10-000-786, Stoelting, Wood Dale, IL). Depiction of infusion cannula placement, fiber optic placement, and virus expression locations are in **Supplemental Fig 1**.

### Drugs

Fentanyl citrate (#22659) obtained from Cayman chemical (Ann Arbor, MI), and deschloroclozapine (#7193) obtained from Tocris Bioscience (Minneapolis, MN), were dissolved in sterile saline (0.9%). The D1 receptor antagonist SCH23390 hydrochloride (#HB1643) obtained from Hello Bio (Princeton, NJ), and the D2 receptor antagonist raclopride (#17422) obtained from Cayman chemical (Ann Arbor, MI), were dissolved in phosphate buffered saline (PBS).

### Conditioned place preference

Conditioned place preference (CPP) was performed as previously described (Montemarano et al. 2025) in a three-chambered rectangular apparatus consisting of a neutral center compartment and two side compartments with different contextual cues, connected by automatic guillotine doors, and equipped with infrared photobeam detectors that track animal position and movement (ENV-3013, MED Associates). On the first day of testing (pre-test), mice underwent a 5 min habituation in the center compartment, after which the doors opened, and they freely explored the entire apparatus for 20 min. The least-preferred chamber was subsequently assigned as the fentanyl-paired chamber (biased design). Mice then underwent 3 consecutive days of conditioning. On conditioning days, mice received an i.p. injection of saline (10 mL/kg) and were confined to the saline-paired compartment for 20 min; 4 h later, they received an i.p. injection of fentanyl (0.2 mg/kg) and were confined to the fentanyl-paired compartment for 20 min. On the fifth day of testing (post-test), after 5 min habituation in the center, they freely explored the entire apparatus for 20 min. To avoid false positives generated by the biased design, mice were considered to have developed fentanyl place preference only if they exhibited both an increase in time in the fentanyl paired context (post-test minus pre-test), and an increased time in the fentanyl-paired relative to saline-paired context (fentanyl-paired minus saline-paired) after conditioning. For the intracranial pharmacology and chemogenetic experiments, these criteria were applied to the post-test under vehicle conditions (i.p. saline for the DREADDs mice and intracranial PBS for the cannula mice). Locomotion was measured as movement counts in Med-PC V Software.

### Intracranial pharmacology

At least 3 days after cannula implantation, mice underwent CPP as described above. Prior to the post-test, mice received intracranial infusions of either the D1R antagonist SCH23390 (0.2 µg / 0.5 µL infused over 1 minute) (See 2009) or PBS. Microinjectors targeting PFC extended 0.5 mm below the guide cannula (-2.2 DV) and for NAc extended 0.1 mm below the guide cannula (-4.1 DV). The infusers were left in place for 5 minutes, then mice started the post-test after an additional 5 minutes. This was repeated four hours later with the opposite treatment (mice that received SCH23390 before the first post-test then received PBS before the second post-test, and vice versa). A separate cohort of mice received intracranial infusions of the D2R antagonist raclopride (1 µg / 0.5 µL infused over 1 minute) (Czachowski et al. 2001) or PBS before the post-test using the same design.

### Fiber photometry

Three weeks after surgery, mice were habituated to a dummy headstage (3g) in their homecage (2-3 days, ∼10 min/day). Changes in GCaMP8s fluorescence were acquired with a TeleFipho Wireless Fiber Photometry system (Amuza, San Diego, CA) and synchronized to behavioral events with TTL pulses via an AD instruments Interface and Lab Chart Lightning software (Colorado Springs, CO). Prior to the first behavioral test, a >10 minute recording was taken to confirm GCaMP expression. A >5 minute recording was taken immediately prior to each CPP session to minimize signal decay during testing. Recordings were taken for pre-test, post-test, and all conditioning sessions (470 nm LED @ ∼30uW excitation power; 500-550 nm emission). After testing completed, mice were perfused and brains processed for immunostaining as described below to confirm viral expression and fiber placement. Custom MATLAB scripts were made for processing & analysis of fiber photometry recordings. To process recordings, TTL timestamps were used to trim recordings from start to end and a double exponential fit of the signal was generated as a pseudoisobestic control for debleaching (Sherathiya et al. 2021). ΔF/F was computed as (F − F_0_)/F_0_, where the baseline fluorescence (F_0_) was obtained from a linear fit of the fluorescence signal over the full 20-minute recording (Lerner et al. 2015). The ΔF/F trace was then z-scored using its mean and standard deviation to generate zΔF/F. Calcium transients were identified using the findpeaks function in MATLAB, using the following criteria: (1) peak height ≥3x median absolute deviation, (2) peak width >300ms, and (3) distance from previous peak ≥500ms. Due to recording errors during conditioning sessions, any mice that did not have all 6 conditioning recordings were excluded from analysis of conditioning data (n=5). For test days, we generated 4 second zΔF/F snippets centered around context entries (±2 sec) from the entire session zΔF/F trace (no additional baseline subtraction). Snippets were filtered such that a mouse must have been inside the saline- or fentanyl-paired context for ≥2 seconds. This cutoff was chosen as it ensures the signals are appropriately aligned to entries (i.e., disallowing multiple entries in one snippet), while still preserving the majority of overall entries (80-85% of all entries are ≥2 seconds). To control for behavioral differences among mice, and ensure an equal number of snippets was included from each mouse, we analyzed the first 15 filtered (≥2s) transitions into each context, which was the minimum number observed across all mice. To compare peri-transition waveforms, we adapted the MATLAB script developed by Jean-Richard-dit-Bressel and colleagues for a bootstrapped confidence interval approach (Jean-Richard-dit-Bressel et al. 2020). For each time point, we computed 95% confidence intervals (α = 0.05) by resampling trials with replacement. Differences between conditions were assessed using bootstrap confidence intervals of the difference waveform (boot_diffCI). Time periods for which the confidence interval did not include zero were considered significant. Because this approach is based on confidence interval estimation rather than hypothesis testing, exact p-values are not reported. No mice were excluded during pre-test or post-test recordings (no technical errors).

### Chemogenetics

At least 14 days after stereotaxic surgery, mice underwent fentanyl CPP as described above, except mice received i.p. injection of saline (or DCZ) 20 min before the post-test. Four hours later, mice were injected with 0.1 mg/kg DCZ (or saline) 20 min before a second post-test (Montemarano et al. 2025). After testing, mice were euthanized, brains removed, and a stereomicroscope equipped with a fluorescence adapter (NightSea, Hatfield PA) was used to confirm GFP-tagged Cre expression in projection targets and mCherry-tagged hM4Di expression in VTA. To validate our chemogenetic approach, a separate cohort of mice received the same stereotaxic surgery to express DREADDs in VTA-NAc or VTA-PFC. After surgical recovery and viral expression (≥14d) mice received an i.p. injection of 0.1 mg/kg DCZ to activate DREADDs, then 0.2 mg/kg fentanyl to induce cFos expression 20 min later. Mice were transcardially perfused ∼1 hour later to capture fos expression as described below.

### Immunostaining

Mice were transcardially perfused with PBS and 4% paraformaldehyde under isoflurane anesthesia. Brains were removed and post-fixed for 24hr, then sectioned to a thickness of 50 µm with a compresstome (Precisionary Instruments, Ashland MA.) Slices from GCaMP brains were washed with PBS and blocked for 30 min in in PBS with 3% normal donkey serum and 0.3% Triton X-100. Slices were incubated at 4°C overnight in blocking buffer containing appropriate primary antibodies (1:1000 chicken anti-GFP, Aves Lab, Tigard, OR, #GFP-1020; 1:1000 rabbit anti-tyrosine hydroxylase [TH], Calbiochem #657012). Slices were washed 3x10 min with PBS, then incubated for 2h in PBS containing secondary antibodies (1:500, Goat anti-Chicken FITC, Aves Lab, #F-1005; Donkey anti-Rabbit Alexa Fluor 594, Jackson Immuno, West Grove, PA, #711-585-152). Slices were washed with PBS 3 x 10 min, counterstained with DAPI, mounted with Fluoromount-G (Southern Biotech, Birmingham, AL), and imaged on a laser-scanning confocal (Leica SP8) or epifluorescence microscope (Nikon Labophot2). For DREADDs validation, slices were washed with PBS and blocking buffer as above then incubated overnight in the appropriate primary antibodies (1:1000 chicken anti-GFP, Aves Lab, Tigard, OR, #GFP-1020; 1:1000 guinea pig anti-RFP, Synaptic Systems, Göttingen, Germany, #390 004; 1:1000 rabbit anti-cFos, Synaptic Systems #226 008). Slices were washed 3x10 min with PBS, incubated for 30 minutes in PBS with 0.5% Triton X-100, then NeuroTrace was added (1:100, Invitrogen #N21479) and slices incubated for an additional 60 min. Slices were then washed 2x5 min then 1x2hr in PBS, then incubated for 2h in PBS containing secondary antibodies (1:500, Goat anti-Chicken FITC, Aves Lab, #F-1005; Donkey anti-guinea pig TRITC, Jackson #706-025-148; Donkey anti-Rabbit Alexa Fluor 647, Jackson #711-605-152). Slices were mounted with Fluoromount-G and imaged on a laser-scanning confocal.

### Cell counting

A subset of wildtype mice received stereotaxic surgery to express GCaMP in VTA-NAc or VTA-PFC neurons as described above, but did not receive a fiber optic cannula to avoid tissue damage artifacts. After >3 weeks, they were perfused and VTA slices were processed for immunostaining of GFP and TH as described above. Images of the right VTA were acquired using a laser-scanning confocal. Three VTA images were pooled per mouse. TH expression was used to outline VTA boundaries, then two blinded investigators independently counted the number of GCaMP-expressing neurons in VTA and how many co-localized with TH to obtain percentage TH-positive GCaMP neurons in VTA. Both investigators’ counts were averaged together to obtain one percentage per image. For cFos validation of DREADDs, images of the entire VTA were acquired using a laser-scanning confocal, pooling 2-4 VTA images per mouse. Boundaries were drawn around the VTA based on landmarks visualized with NeuroTrace, then two investigators independently obtained percentage cFos-positive mCherry neurons in VTA as above.

### RNA Isolation & Sequencing

Mice were euthanized by cervical dislocation, and brains were rapidly removed and chilled in ice cold PBS. Cold brains were cut into 1 mm coronal sections using an aluminum brain matrix, and tissue punches containing total VTA were collected with 14-gauge needle, then snap-frozen on dry ice and stored at -80 until processing.

Polyribosomes were immunoprecipitated as described previously (Fox et al. 2020, 2023). Tissue punches from RiboTag mice (3 mice pooled per sample) were homogenized by douncing in 1 mL homogenization buffer, and 50µL supernatant saved as input. Remaining supernatant was incubated with 5 µl anti-HA antibody (BioLegend #901515) at 4°C overnight with constant rotation. Samples were then incubated with 400 uL of protein A/G magnetic beads (Thermo Scientific #88803) overnight at 4°C with constant rotation. Beads were washed in a magnetic rack with high-salt buffer. RNA was extracted using Trizol (Invitrogen) and the RNeasy Mini Kit with a DNAse step (Qiagen) from both immunoprecipitated and input samples by adding lysis buffer supplemented with beta mercaptoethanol and following the manufacturer’s instructions (3-5 samples/sex/projection, 45 total mice). RNA quality and concentration were assessed using an Agilent BioAnalyzer, and cDNA libraries were prepared by the PSUCOM Genome Sciences Core using the SMARTer Stranded Total RNA-Seq Kit v3 – Pico Input Mammalian (Takara Bio, San Jose, CA) according to the manufacturer’s instructions. Libraries were quantified by qPCR prior to sequencing on an Illumina NovaSeq 6000 (50 base pair paired-end reads; average of ∼70 million reads per sample). Raw data were demultiplexed using bcl2fastq (Illumina), and adapter trimming was performed after UMI extraction with UMI-tools (Smith et al. 2017). Sequencing reads were aligned to the mouse reference genome (GRCm39, Ensembl release 2.7.1a) using STAR (Dobin et al. 2013) with a maximum of two mismatches allowed per read. Gene-level counts were obtained using featureCounts (Liao et al. 2014), with multi-mapping reads included, reads assigned to multiple overlapping features counted fractionally, and the feature with the largest overlap prioritized. We used limma (lmfit) (Ritchie et al. 2015) in R (v4.5.2) to normalize counts (log2CPM) and determine differential expression between projection neurons vs total VTA with post hoc contrasts (makeContrasts). To asses cell-type enrichment across projections, we used the mean_ratios function from DeconvoBuddies (Huuki-Myers et al. 2025) in conjunction with our previously published snRNAseq dataset of mouse VTA (Fox et al. 2024). The top 50 markers per cluster with a MeanRatio >1 were selected. Overlap between differentially expressed genes and cluster markers was evaluated using Fisher’s exact test. For each projection, the proportion of differentially expressed genes mapping to each marker-defined cell class was calculated as the number of overlapping genes divided by the total number of differentially expressed genes overlapping any cluster marker.

### Data analysis

Except where noted, data were analyzed with GraphPad Prism (v10), and were collapsed across sex in the absence of significant sex effects or interactions. Mice with off-target viral or cannula placements, or not exhibiting fentanyl place preference, were excluded (10 NAc cannula mice, 16 PFC cannula mice, 5 GCaMP mice, and 10 DREADDs mice). For intracranial pharmacology experiments, data were analyzed with paired t-tests. For calcium activity during drug exposure, we only included mice with all 6 conditioning recordings (n=14 out of 19 total mice), and data were analyzed with paired t-tests. Peri-event signal was analyzed with bootstrapped 95% confidence intervals in MATLAB (Jean-Richard-dit-Bressel et al. 2020), and mean signals within a specific time window (analogous to AUC) were analyzed with paired t-tests. For chemogenetic experiments, data were analyzed with two-way repeated measures ANOVA with factors of virus and agonist, and significant interactions were followed up with Sidak’s post hoc.

## RESULTS

### Fentanyl context-seeking is attenuated by D1R blockade in PFC but not NAc

Dopaminergic mechanisms regulating drug-context associations vary across misused drugs. It remains unknown which dopamine receptors promote fentanyl context-seeking. To address this, we locally blocked D1Rs in PFC or NAc during fentanyl CPP expression. Using a counterbalanced, within-subject design, mice underwent two post-tests 4 hours apart, each preceded by an intracranial infusion of either vehicle or the D1 receptor antagonist SCH23390 (Timeline in **Fig 1A**, example cannula placements in **Fig 1B, F**). We found that intra-NAc infusion of SCH23390 had no effect on fentanyl place preference compared to vehicle (**Fig 1C-D**). The absence of behavioral effect was not due to an ineffective dose, as intra-NAc infusion of SCH23390 did reduce locomotion (**Fig 1E**; t_17_=2.9, p=0.01). In contrast, intra-PFC infusion of SCH23390 reduced time spent in the fentanyl-paired side (**Fig 1G**; t_17_=2.17, p=0.045) and preference for the fentanyl over saline side (**Fig 1H**; t_17_=2.35, p=0.031) relative to vehicle. Intra-PFC infusion of SCH23390 did not impact locomotion (**Fig 1I**).

**Figure 1.**
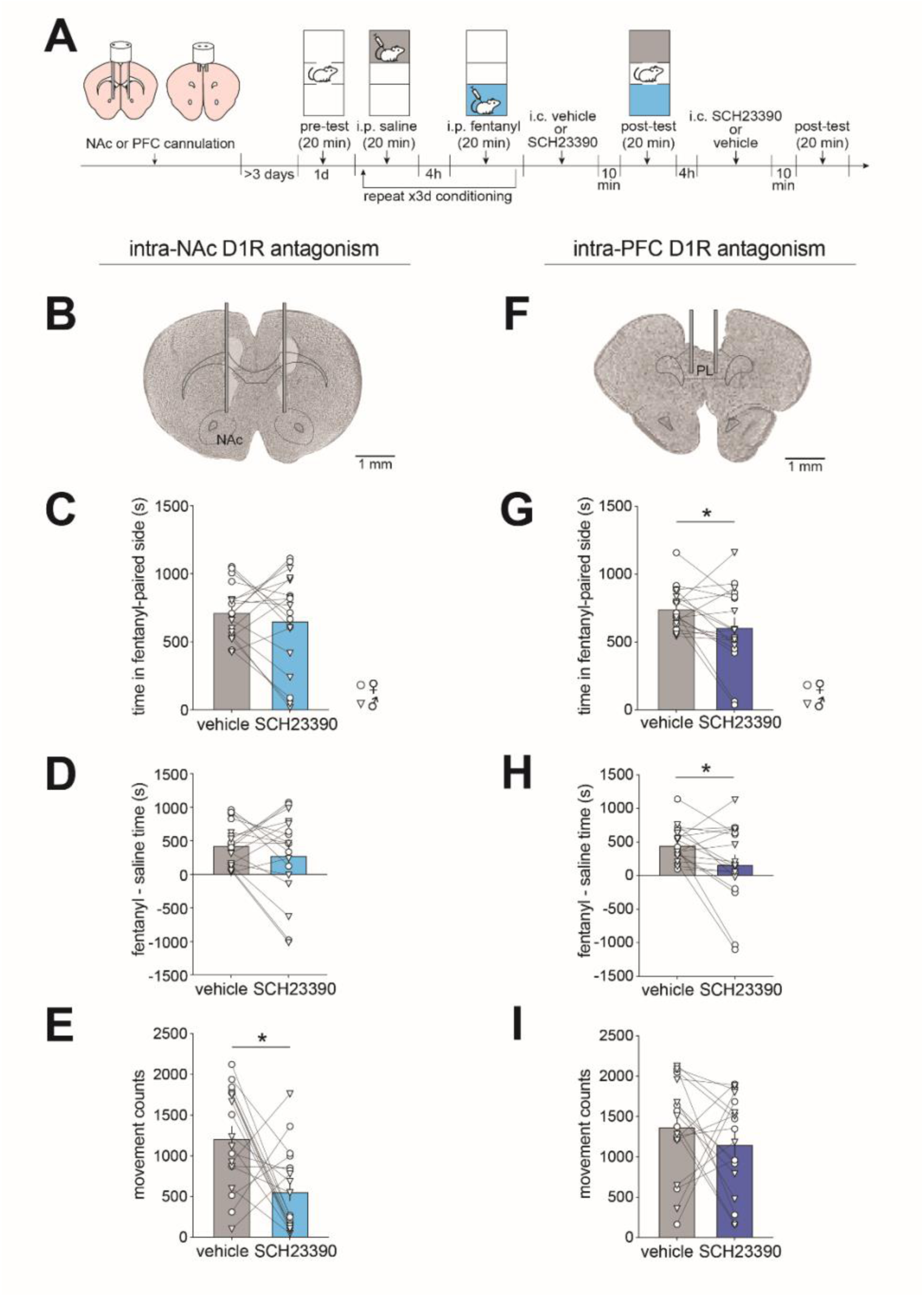
D1R antagonism in PFC, not NAc, attenuates fentanyl context-seeking. (**A**) Timeline for intracranial SCH23390 experiments. Mice received a bilateral infusion cannula in either NAc or PFC, and underwent fentanyl CPP at least 3 days later. Ten minutes prior to each post-test, mice received intracranial SCH23390 (0.2 µg / 0.5 µL infused over 1 minute) or vehicle in a counterbalanced repeated-measures design. (**B**) Representative image of NAc cannula placement. Intra-NAc SCH23390 had no effect on (**C**) time in the fentanyl-paired context, or (**D**) preference for the fentanyl- over saline-paired context. (**E**) Intra-NAc SCH23390 reduced movement counts relative to vehicle (n=18, *, p=0.01). (**F**) Representative image of PFC cannula placement (PL, prelimbic cortex). Intra-PFC SCH23390 reduced (**G**) time in the fentanyl-paired context (n=18, *, p=0.045) and (**H**) preference for the fentanyl- over saline-paired context (*, p=0.031) relative to vehicle. (**I**) Intra-PFC SCH23390 had no effect on movement counts. Data are presented as mean±SEM with individual mice overlaid. Circles represent data points from females, and triangles from males. Depictions of all cannula placements available in **Supplemental Figure 1**.

### Fentanyl context-seeking is attenuated by D2R blockade in NAc but not PFC

Given the region-dependent effects of D1 receptor blockade on fentanyl CPP, we next investigated D2 receptors. We administered the D2 receptor antagonist raclopride in either PFC or NAc immediately prior to the post-test (**Fig 2A**). In contrast to the D1R antagonist, intra-NAc infusion of raclopride reduced time spent in the fentanyl-paired side (**Fig 2B**; t_15_=2.95, p=0.0099) and preference for the fentanyl over saline side (**Fig 2C**; t_15_=2.71, p=0.016) compared to vehicle. Intra-NAc raclopride also reduced locomotion (**Fig 2D**; t_15_=3.04, p=0.0083). In contrast, intra-PFC raclopride had no effect on fentanyl place preference (**Fig 2E-F**) or locomotion (**Fig 2G**).

**Figure 2.**
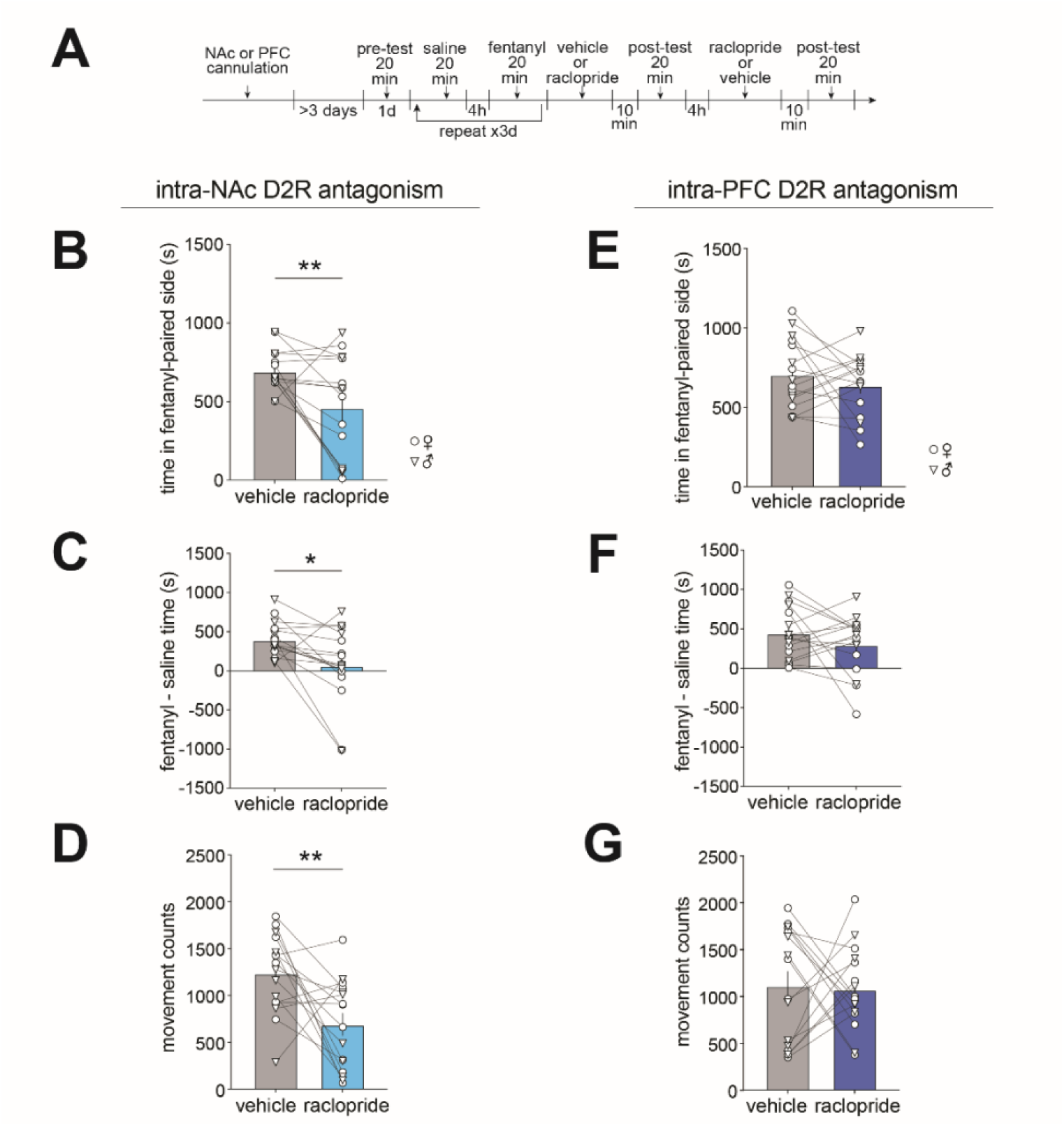
D2R antagonism in NAc, not PFC, attenuates fentanyl context-seeking. (**A**) Timeline for intracranial raclopride experiments (1 µg raclopride / 0.5 µL infused over 1 minute). Intra-NAc raclopride reduced (**B**) time in the fentanyl-paired context (n=16, **, p=0.0099), (**C**) preference for the fentanyl- over saline-paired context (*, p=0.0116) and (**D**) movement counts (**, p=0.0083) relative to vehicle. Intra-PFC raclopride had no effect on (**E**) time in the fentanyl-paired context, (**F**) preference for the fentanyl- over saline-paired context, or (**G**) movement counts, relative to vehicle (n=16). Data are presented as mean±SEM with individual mice overlaid. Circles represent data points from females, and triangles from males.

### Both VTA-NAc and VTA-PFC exhibit elevated calcium activity during fentanyl conditioning

Since we found a role for PFC and NAc dopamine receptors in fentanyl CPP expression, we next investigated upstream VTA signaling directed at each of these target regions. While there is support for dopamine-independent mechanisms of opioid CPP (Hnasko et al. 2005), whether these mechanisms rely on VTA is unknown. Therefore, we targeted VTA-NAc and VTA-PFC projection neurons in a cell type-agnostic manner to investigate their respective roles in fentanyl CPP. We used fiber photometry to measure calcium activity of VTA-NAc and VTA-PFC during fentanyl CPP (**Fig 3A**). First, we established the cell-types captured by our viral approach. We found heterogeneity in projection-specific cell types expressing GCaMP, with 32.5% of VTA-NAc GCaMP-expressing neurons, and 28.4% of VTA-PFC GCaMP-expressing neurons, colocalizing with TH (**Fig 3B**; n=5 mice per projection target group x 3 images per mouse; VTA-NAc vs VTA-PFC t-test, t_8_=1.22, p=0.26). We corroborated findings at the protein level with mRNA using the RiboTag mouse. We sequenced ribosome-associated mRNA from projection neurons alongside total VTA, and used enrichment of cell-type marker genes to estimate the proportion of dopamine, glutamate, and GABA neurons. Similar to IHC, ∼20% of each projection was enriched for dopamine markers, and the remaining 80% was evenly split between glutamatergic and GABAergic markers (**Supplemental File 1**).

**Figure 3.**
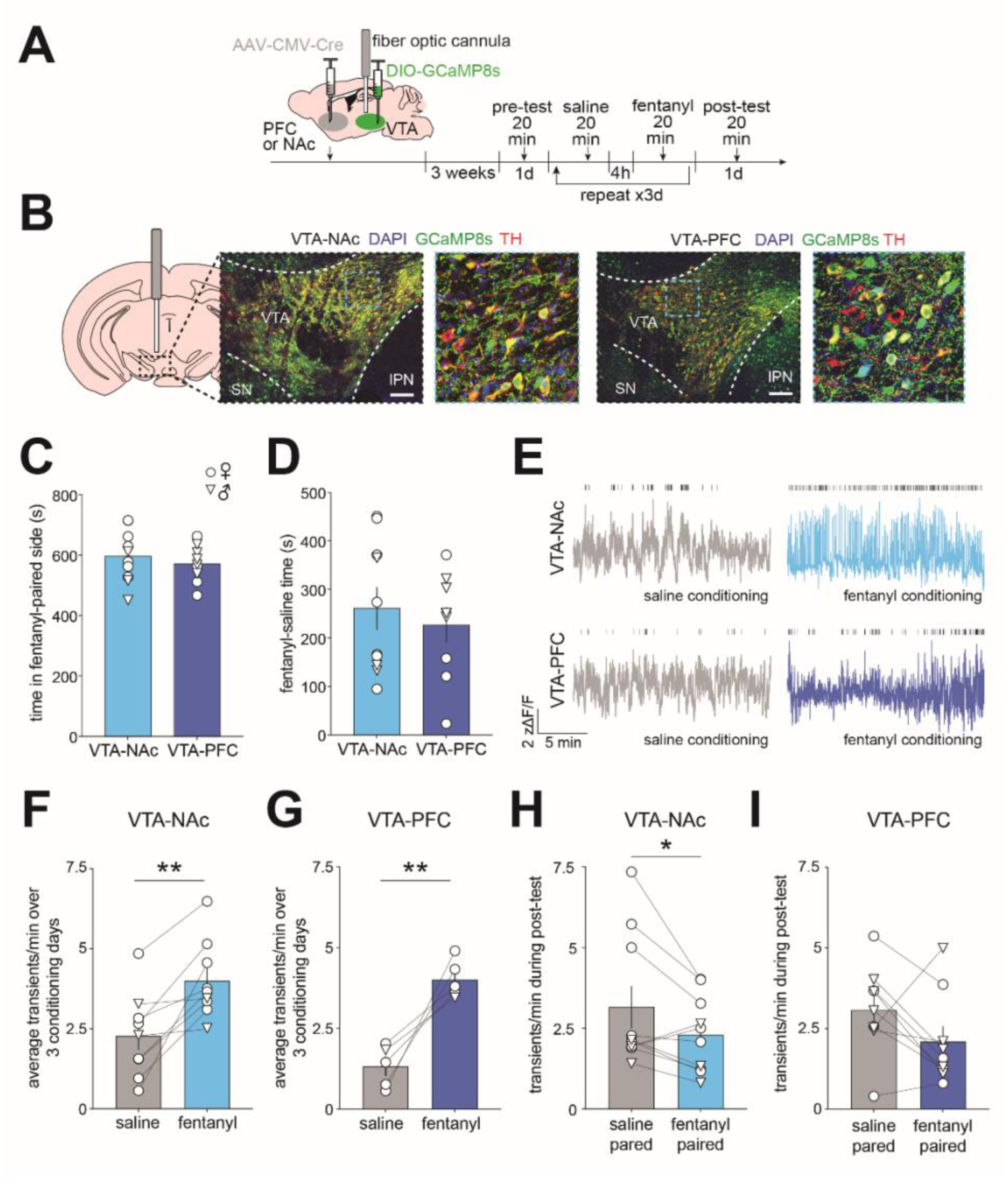
VTA-NAc and VTA-PFC have elevated calcium activity in response to fentanyl exposure. (**A**) Timeline for projection-specific VTA GCaMP fiber photometry experiments. Mice received retrograde Cre in either the PFC or NAc, and Cre-dependent GCaMP8s in VTA, with a fiber optic cannula implanted directly above VTA. Three weeks later, mice underwent fentanyl CPP and VTA GCaMP signal was recorded during all sessions. (**B**) Representative images depicting GCaMP expression in VTA-NAc and VTA-PFC with immunostaining for TH (SN, substantia nigra, IPN, interpeduncular nucleus). (**C**) VTA-NAc and VTA-PFC GCaMP groups did not differ in time spent in fentanyl-paired side during the post-test, or (**D**) preference for the fentanyl- over saline-paired context (VTA-NAc n=10, VTA-PFC n=9). (**E**) Representative calcium traces from VTA-NAc (top) and VTA-PFC (bottom) during saline conditioning (left) and fentanyl conditioning (right). Ticks above the trace indicate identified calcium transients. (**F**) In VTA-NAc, fentanyl increases transient frequency (n=9, average over 3 conditioning days, **, p=0.0025) (**G**) In VTA-PFC, fentanyl increases transient frequency (n=5, average over 3 conditioning days, **, p=0.0074). (**H**) During the post-test, VTA-NAc has reduced transient frequency when the mouse is in the fentanyl- compared to saline-paired context (n=10; *, p=0.046) (**I**) No difference in VTA-PFC transient frequency in saline-paired vs fentanyl-paired context (n=9; p=0.1). Data are presented as mean±SEM with individual mice overlaid. Circles represent data points from females, and triangles from males. All viral and fiber optic placements are available in **Supplemental Figure 1**.

To determine the involvement of VTA-NAc and VTA-PFC in fentanyl CPP, we first ensured no behavioral differences across the groups. Both VTA-NAc and VTA-PFC GCaMP mice spent a similar amount of time in the fentanyl-paired context during the post-test (**Fig 3C**; VTA-NAc vs VTA-PFC t-test, t_17_=0.72, p=0.48) and exhibited similar preference for the fentanyl-paired over the saline-paired context (**Fig 3D**; VTA-NAc vs VTA-PFC t-test, t_17_=0.59, p=0.56). To examine activity of VTA-NAc and VTA-PFC neurons in response to fentanyl we analyzed photometry data during the conditioning sessions (**Fig 3E**). There were no main effects of conditioning day, so data were averaged across days and analyzed with paired t-test. For both VTA-NAc and VTA-PFC neurons, we found a greater number of calcium transients during fentanyl conditioning relative to saline (VTA-NAc: **Fig 3F**, t_8_=4.32, p=0.0025; VTA-PFC: **Fig 3G**, t_4_=5.018, p=0.0074). We then analyzed transient frequency during the post-test and found VTA-NAc had reduced transient frequency when the mouse was in the fentanyl-paired context compared to the saline-paired context (**Fig 3H**, t_9_=2.32, p=0.046) while VTA-PFC had no significant difference in frequency in the fentanyl- vs saline-paired context (**Fig 3I**, t_8_=1.84, p=0.1).

### Both VTA-NAc and VTA-PFC exhibit elevated calcium activity during entries to the fentanyl-paired context

Next, we asked if VTA-NAc and VTA-PFC were differentially engaged during entry into the fentanyl-paired context on test days, as others have shown VTA calcium increases during entry to a cocaine-paired context (Calipari et al. 2017). First, to ensure signals reflected entry into a single context, we restricted analysis to entries where mice spent ≥2s in the fentanyl or saline-paired chamber. Second, to control for behavioral differences and to ensure equal contribution from all mice, we analyzed the first 15 transitions into each side of the chamber for each mouse (the lowest common denominator). Using bootstrapped 95% confidence intervals to determine where signals differed from zero, and from each other (Jean-Richard-dit-Bressel et al. 2020), we found increased VTA-NAc activity preceding entries to both the saline- and fentanyl-paired contexts, but the activity was greater in the one second immediately preceding fentanyl-paired entries compared to saline-paired entries (**Fig 4A**). In contrast, VTA-PFC neurons exhibited increased activity only preceding fentanyl-paired, but not saline-paired entries (**Fig 4C**). For both projections, increased calcium activity during entries to the fentanyl-paired context did not reflect exits from the saline-paired context, since there were no differences in the signal when aligned to region exits (**Fig 4A,C**). The increased activity preceding fentanyl-paired context entry was consistent across the first 15 filtered entries for both projections (**Fig 4B,D**), but not preceding saline-paired entries (**Supplemental Fig 2**). To ensure these effects were consistent within animals, for each animal we calculated the mean zΔF/F over the 1 second immediately preceding region entries (-1 to 0 seconds from entry) and used paired t-tests to compare between saline- and fentanyl-paired entries. During the pre-test, there were no differences in mean zΔF/F preceding region entries for both VTA-NAc (**Fig 4E**) and VTA-PFC (**Fig 4G**), ensuring that the increased activity observed during the post-test is due to context associations and not pre-existing biases. After conditioning, like what we observed with bootstrapped confidence intervals, there was greater mean zΔF/F in the one second preceding fentanyl-paired entries compared to saline-paired entries for both VTA-NAc (**Fig 4F**, t_9_=2.39, p=0.041) and VTA-PFC (**Fig 4H**, t_8_=2.53, p=0.035).

**Figure 4.**
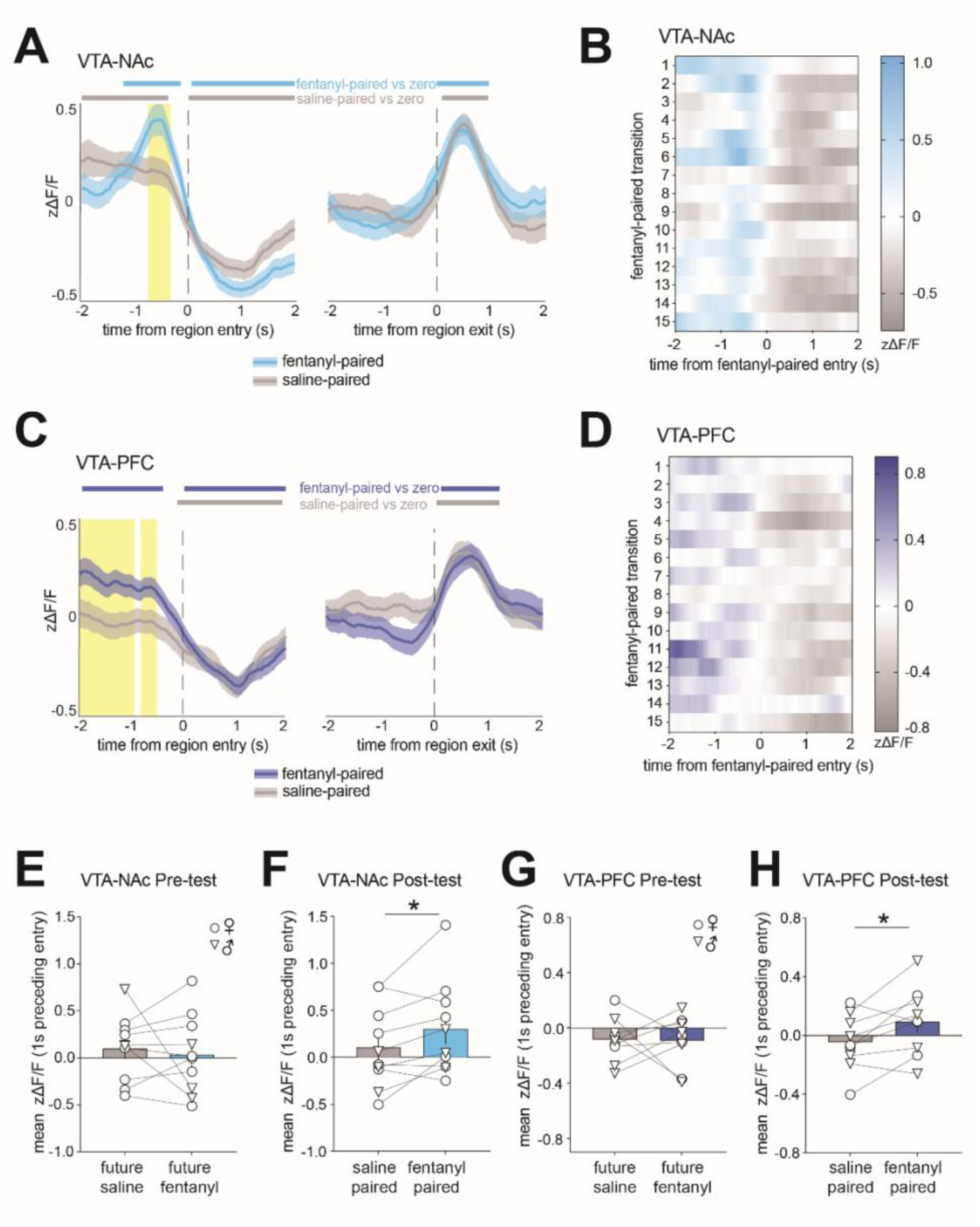
VTA-NAc and VTA-PFC have elevated calcium activity during entries to the fentanyl-paired context. (**A**) Average zΔF/F for all VTA-NAc mice for 4 seconds surrounding the first 15 filtered (≥2s) entries (left) or exits (right) to the saline-paired (gray trace) or fentanyl-paired (blue trace) context. Vertical dotted line represents the time of entry/exit. Using bootstrapped 95% confidence intervals, gray and blue horizontal bars represent where the gray and blue traces (saline-paired and fentanyl-paired, respectively) significantly differ from 0, and yellow shaded area represents where the two traces significantly differ from one another. (**B**) Heatmap of VTA-NAc activity for the 4 seconds surrounding the first 15 filtered entries into the fentanyl-paired region. Color represents zΔF/F averaged across all VTA-NAc mice. (**C-D**) Same as A-B, but for VTA-PFC. (**E**) During the pre-test for VTA-NAc there is no difference in mean zΔF/F over the 1 second preceding entry to either context. (**F**) During the post-test for VTA-NAc there is greater calcium activity in the 1 second preceding entries to fentanyl-paired vs saline-paired context (n=10; *, p=0.04). (**G-H**) Same as E-F, but for VTA-PFC (n=9, *, p=0.035). Data are presented as mean±SEM with individual mice overlaid. Circles represent data points from females, and triangles from males.

### Chemogenetic inhibition of VTA-NAc, not VTA-PFC, attenuates fentanyl context-seeking

Next, we used chemogenetic inhibition to determine whether activity in VTA-NAc and VTA-PFC functionally support fentanyl context-seeking. We expressed the inhibitory DREADD hM4Di, or an mCherry control, in either VTA-NAc or VTA-PFC (Timeline in **Fig 5A**). To validate our approach, a subset of mice were injected with DCZ, then with fentanyl to induce cFos expression (**Fig 5B-C**). There was ∼50% less cFos in hM4Di neurons relative to mCherry neurons in both projection populations (**Fig 5D**. hM4Di vs mCherry: VTA-NAc: 27.47±2.29 vs 15.05±2.03 cFos+ cells; VTA-PFC: 20.17±1.23 vs 9.55±0.93 cFos+ cells; VTA-NAc vs VTA-PFC p=0.42). The remaining mice underwent CPP as before. Using a counterbalanced, within-subject design, mice underwent two post-tests 4 hours apart, each preceded by an injection of either DCZ or vehicle (**Fig 5A**). Chemogenetic inhibition of VTA-NAc reduced time spent in the fentanyl-paired context (**Fig 5E**, vehicle vs DCZ: hM4Di p=0.036, mCherry p=0.81, Sidak’s post hoc) as well as preference for fentanyl- over saline-paired context (**Fig 5F**; vehicle vs DCZ: hM4Di p=0.018, mCherry p=0.91, Sidak’s). This effect was not due to reduced locomotion (**Fig 5G**). In contrast, chemogenetic inhibition of VTA-PFC did not significantly alter fentanyl place preference (**Fig 5H-I**), but did reduce movement counts (**Fig 5J**, vehicle vs DCZ: hM4Di p=0.037, mCherry p=0.32, Sidak’s post hoc,).

**Figure 5.**
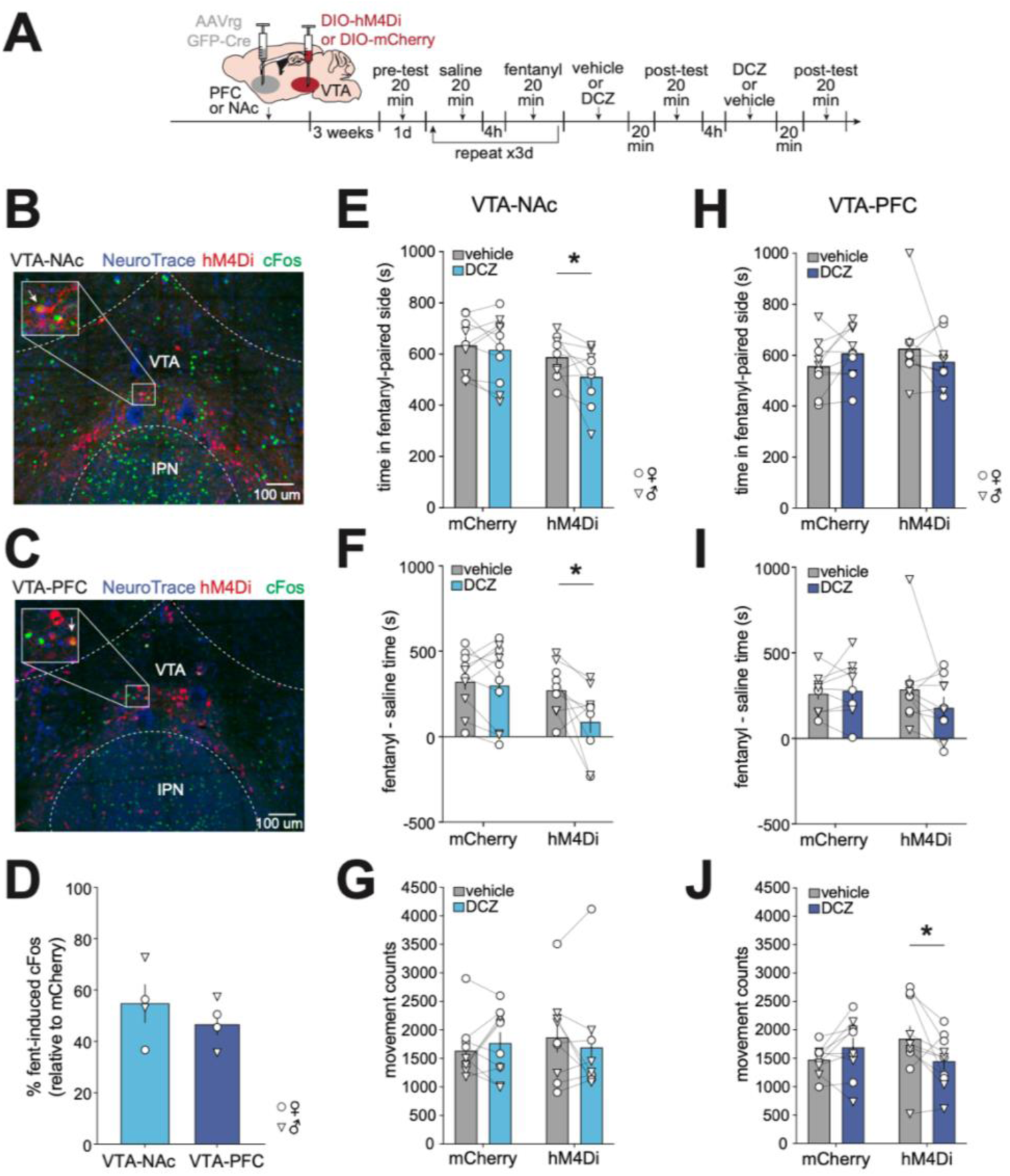
Chemogenetic inhibition of VTA-NAc, not VTA-PFC, attenuates fentanyl context-seeking. (**A**) Timeline for chemogenetic experiments. Mice received retrograde-Cre in either PFC or NAc, and Cre-dependent hM4Di, or Cre-dependent mCherry, in VTA. Twenty minutes prior to each post-test, mice received DCZ (0.1 mg/kg i.p.) or vehicle in a counterbalanced repeated-measures design. Representative images of hM4Di and cFos expression in (**B**) VTA-NAc and (**C**) VTA-PFC. (**D**) Both VTA-NAc and VTA-PFC exhibit similar reduction in fentanyl-induced cFos expression in hM4Di relative to mCherry neurons (n=3-4 mice/virus/projection). (**E**) Time in fentanyl-paired context (*, p=0.036), (**F**) preference for fentanyl- over saline-paired context (*, p=0.018), and (**G**) total movement counts after treatment with vehicle (gray) and DCZ (light blue) in mice expressing mCherry (n=10) or hM4Di (n=9) in VTA-NAc. (**H**) Time in fentanyl-paired context, (**I**) preference for fentanyl- over saline-paired context, and (**J**) total movement counts (*, p=0.037) after treatment with vehicle (gray) and DCZ (dark blue) in mice expressing mCherry (n=10) or hM4Di (n=10) in VTA-PFC. Data are presented as mean±SEM with individual mice overlaid. Circles represent data points from females, and triangles from males. All viral placements are depicted in **Supplemental Figure 1.**

## DISCUSSION

In this study we investigated mesolimbic and mesocortical regulation of fentanyl-context associations. We showed a role for dopaminergic signaling in both PFC and NAc, where fentanyl context-seeking is supported by activity at D1 receptors in PFC, and activity at D2 receptors in NAc. To investigate potential roles for other neurotransmitters in addition to dopamine, we targeted VTA-NAc and VTA-PFC neurons in a cell type-agnostic manner. Despite increased calcium activity in both VTA-NAc and VTA-PFC neurons during fentanyl CPP, only chemogenetic inhibition of VTA-NAc, not VTA-PFC neurons, attenuated the expression of fentanyl CPP. Together these data demonstrate that fentanyl context-seeking engages VTA projections to both NAc and PFC, but only VTA-NAc neurons directly modulate the behavior.

### Divergent dopaminergic signaling mechanisms in NAc and PFC during fentanyl context-seeking

To address gaps in knowledge regarding dopaminergic signaling mechanisms that promote fentanyl CPP, we administered local dopamine receptor antagonists during the post-test. This approach allowed us to understand receptor mechanisms in the expression of place preference without also influencing learning of drug-context associations. In PFC, we found an effect of D1 but not D2 receptor antagonism. This is somewhat unsurprising, as D1R is more abundant than D2R in PFC (Santana et al. 2009), though there is some overlap between these populations, with ∼14% of D1R+ PFC neurons co-expressing D2R (Wei et al. 2018). Our findings with fentanyl also resemble those with cocaine, as blocking D1R but not D2R in PFC reduces the expression of cocaine CPP (Shinohara et al. 2017). While our study is the first to demonstrate this in fentanyl CPP, others have shown similar mechanism in other forms of opioid seeking, as PFC D1R blockade attenuates cue-induced heroin seeking after self-administration (See 2009). Since CPP reflects a form of contextual drug cue seeking, these findings suggest that dopaminergic mechanisms of drug-cue recognition in PFC may be consistent across drug classes and behavioral paradigms.

In NAc, we found that blocking D2R, but not D1R, inhibited the expression of fentanyl CPP. Both D1 and D2 receptors are equally abundant on GABAergic medium spiny neurons (MSNs) in NAc with little to no overlap in their expression patterns (Reiner et al. 2024). While D1-MSNs are classically known for promoting motivation and reward seeking, there is evidence to suggest these processes can also be mediated by D2-MSNs (Soares-Cunha et al. 2016; Gallo et al. 2018). Furthermore D2R, unlike D1R, is expressed on other cell types in the striatum besides MSNs, such as dopaminergic afferents (Sesack et al. 1998; Benoit-Marand et al. 2001) and cholinergic interneurons (Yan et al. 1997; Maurice et al. 2004). As such, D2R inhibition likely has a more widespread effect on intra-NAc signaling as compared to D1R inhibition, which could further explain the behavioral differences resulting from these antagonists. An additional effect of intra-NAc dopamine receptor blockade is that both antagonists reduced locomotion during the post-test. This effect has been previously reported, with intra-NAc SCH23390 or raclopride reducing locomotion in a CPP apparatus (Young et al. 2014). Importantly, we believe the observed reduction in fentanyl-context preference following intra-NAc D2R blockade is likely not driven by reduced locomotion, as intra-NAc D1R blockade reduced locomotion without affecting preference for the fentanyl-paired context.

### VTA-NAc, not VTA-PFC projection neurons functionally support fentanyl context-seeking

Having determined a role for dopaminergic signaling in both NAc and PFC during the expression of fentanyl CPP, we next investigated activity of upstream VTA neurons that project to each of these regions. Prior research suggests that opioid CPP is not wholly dependent on dopaminergic mechanisms. Dopamine deficient mice (Hnasko et al. 2005) and D1R-knockout mice (Ting-A-Kee et al. 2013) can both acquire morphine CPP, but the extent to which these effects are mediated by the VTA is unknown. As such, we broadened our investigation to include other neuron types in VTA-NAc and VTA-PFC pathways besides just dopamine neurons. To achieve this, we opted for a projection-specific, but cell type-agnostic AAV approach that enriches for non-dopamine neurons, as midbrain dopamine neurons are not efficiently captured by most AAV serotypes (Tervo et al. 2016). Indeed, our immunolabelling revealed only ∼30% of GCaMP-expressing neurons were TH-positive in both projections. This effect was further corroborated by our Ribotag RNAseq, showing about 20% of each projection expressed dopamine markers, and the remaining 80% glutamate and GABA markers. At both RNA and protein level, our AAV strategy captured a lower proportion of dopamine neurons as compared to other approaches. For example, retrograde tracer Fluorogold reveals ∼70% of mouse VTA projection neurons are TH+ (Taylor et al. 2014), while Cre-expressing canine adenoviruses show ∼60% VTA-NAc are TH+ vs 40% VTA-PFC (Beier et al. 2015). We suspect our low proportion of TH-expression in VTA projection neurons reflects AAV transduction inefficiency and is not a true reflection of the endogenous proportion of dopamine neurons in these projections. However, by enriching for multiple cell types in these pathways, we reveal ways in which diverse VTA cell types may support fentanyl CPP through their projections to other brain regions.

In the present study we found both VTA-NAc and VTA-PFC had increased calcium activity during fentanyl exposure, consistent with classic opioid-mediated disinhibition in the VTA. While these signals may also reflect learning of the drug-context association during conditioning, it is impossible to separate this from the effects of drug exposure on neuronal activity. Others have shown increased calcium activity in VTA neurons and their downstream projections to NAc during entries to a previously cocaine-paired context (Calipari et al. 2017). We thus aligned our calcium signal to context transitions to look at activity during entries to the fentanyl-paired context while animals are drug free. We found similar effects in both VTA-NAc and VTA-PFC neurons, where both had greater calcium activity immediately preceding fentanyl-paired entries compared to saline-paired entries. To determine whether this activity promotes fentanyl CPP or is a consequence of CPP, we chemogenetically inhibited either projection during the post-test. Surprisingly, we found only chemogenetic inhibition of VTA-NAc, but not VTA-PFC neurons attenuated fentanyl CPP expression. Collectively, this suggests that while VTA-PFC neurons are engaged during fentanyl context-seeking, their causal role is more limited relative to VTA-NAc neurons.

Beyond dopamine, VTA glutamate and GABA projection neurons have known roles in motivated behavior and reinforcement learning (Root et al. 2014, 2020; Wang et al. 2015; Yoo et al. 2016; Zell et al. 2020; Warlow et al. 2024). VTA glutamate neurons can modulate oxycodone seeking through their local connections to mesolimbic dopamine neurons (McGovern et al. 2023), and both VTA glutamate and GABA neurons undergo diverse transcriptional adaptations after fentanyl self-administration (Fox et al. 2024). Downstream, blocking GABA and glutamate receptors in NAc can attenuate CPP (Popik and Kolasiewicz 1999; Maguire et al. 2014; Qi et al. 2015), however these effects cannot necessarily be ascribed to VTA neurotransmission. Here, our data that overrepresent non-dopamine neurons indicate some of these glutamatergic and GABAergic effects may be driven by the VTA. While VTA-NAc glutamate opposes cocaine seeking (Barbano et al. 2024) this may not extend to fentanyl. However, it cannot be ruled out that chemogenetic inhibition of a subset of VTA-NAc dopamine neurons was sufficient to attenuate fentanyl CPP expression, especially considering the importance of downstream NAc dopamine receptors for this behavior. Additional studies should investigate the role for neurotransmitter-defined cell types across a variety of opioid administration paradigms, as our data indicate VTA-NAc neurons as a whole are important for the expression of fentanyl-context associations.

### Conclusions

Here we demonstrate that projection-specific VTA neurons drive the expression of fentanyl-context associations through diverse signaling, providing a unique viewpoint of VTA representations of opioid behaviors. These experiments lay the foundation for further investigation into cell type-specific encoding of opioid-associated cues in the VTA. These findings provide necessary insight to the circuit-level neuroadaptations that promote fentanyl context-seeking, thus improving our understanding of potential neural mechanisms underlying OUD.

## Acknowledgements

The authors thank the Penn State College of Medicine Genome Sciences Core (RRID:SCR_021123) and Advanced Light Microscopy Core (RRID:SCR_022526). A previous version of this manuscript was deposited to bioRxiv.

## Author Contributions

AM and MEF designed the experiments. AM and FAA performed surgeries. AM, HS, LBM, LDF, and SP performed behavior experiments. MEF performed RNA isolation. AM and KD counted cells. AM and MEF performed data analysis. AM and MEF wrote the manuscript and prepared the figures with input from all authors.

## Funding

This work was supported by NIH grant DA058661 awarded to MEF.

## Competing Interests

The authors have nothing to disclose.

## Data and code availability

Code is available at github.com/mfoxlab. RNAseq data have been deposited at GEO and are publicly available as of the date of publication. Other data are available upon reasonable request addressed to the corresponding author.

## Notes

### Competing Interest Statement

The authors have declared no competing interest.

### Summary of Updates

Supplemental figures were added, and minor text edits were made to address reviewer feedback.

