## Supplemental Figures for "Mesolimbic and mesocortical pathways differentially support fentanyl-context associations"

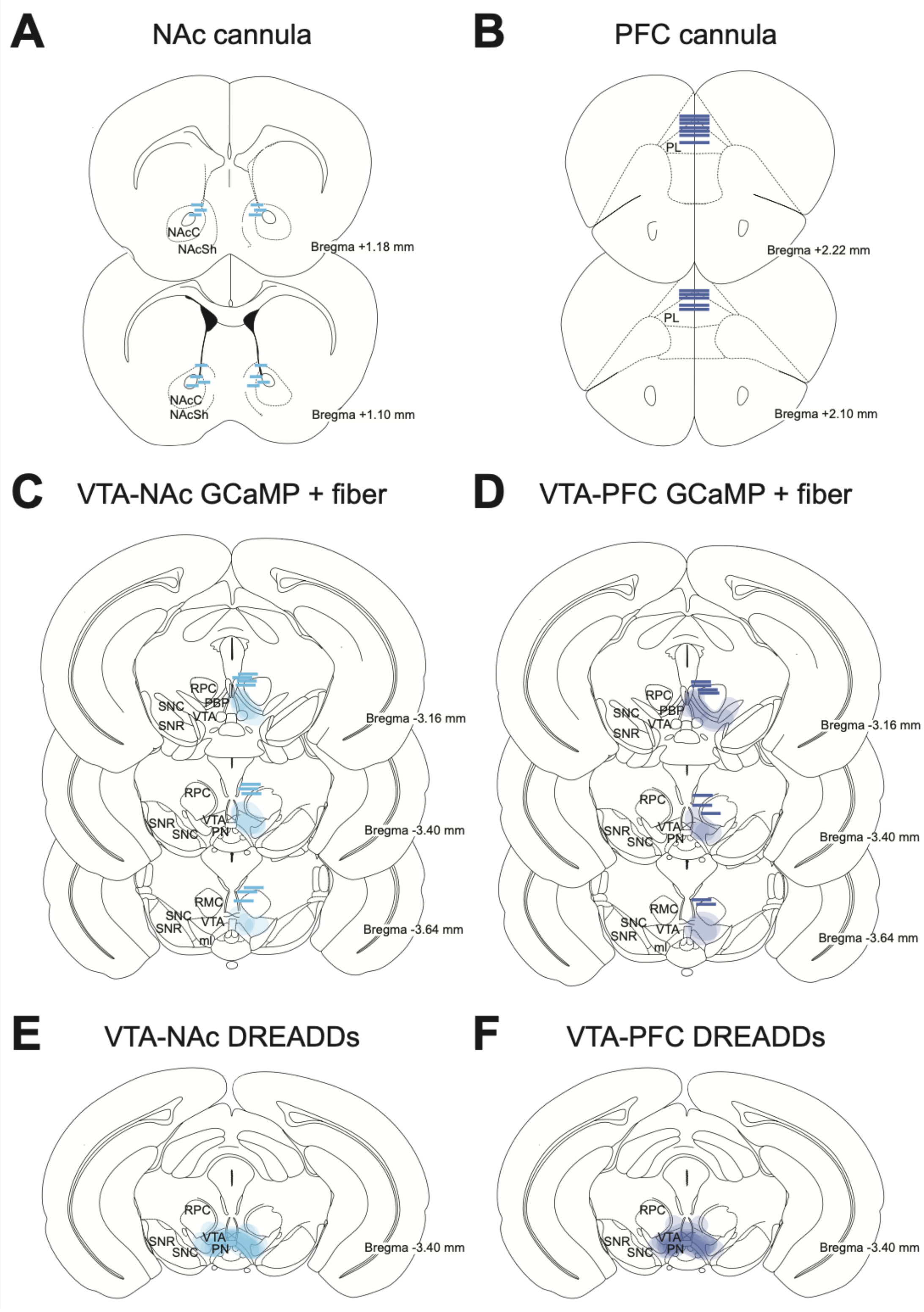


**Supplemental Figure 1.** Verification of cannula & virus locations. (**A-B**) Blue lines depict termination of NAc or PFC infusion cannulas. NAcC, nucleus accumbens core. NAcSh, nucleus accumbens shell. PL, prelimbic cortex. (C-D) Blue lines depict termination of fiber optic, and blue shaded area depicts GCaMP expression in VTA-NAc or VTA-PFC neurons. (E-F) Blue shaded area depicts hM4Di or mCherry expression in VTA-NAc or VTA-PFC neurons. Darker shaded area indicates more animals showed expression in that area. RMC, red nucleus, magnocellular. RPC, red nucleus, parvocellular. SNC, substantia nigra, pars compacta. SNR, substantia nigra, pars reticulata. PBP, parabrachial pigmented nucleus of VTA. PN, paranigral nucleus of VTA. ml, medial lemniscus.

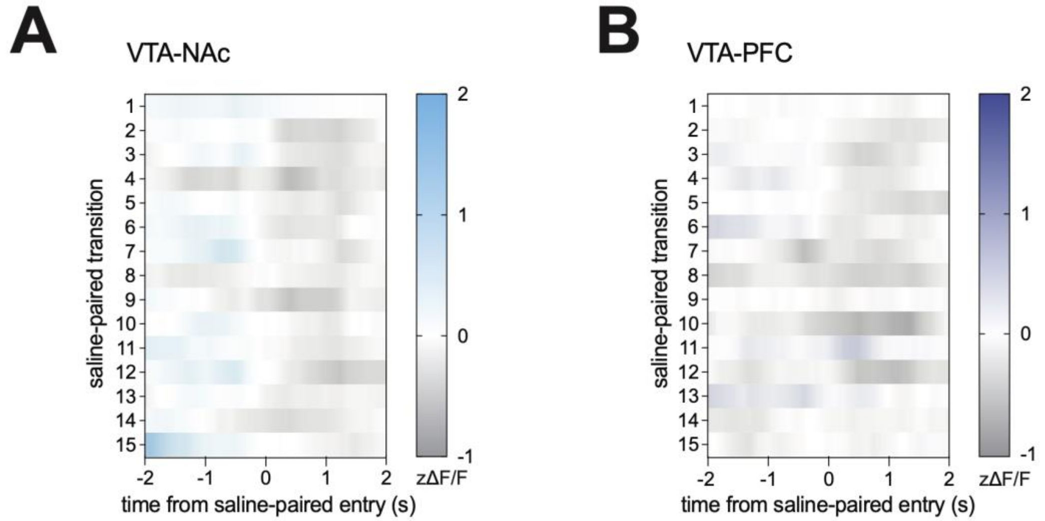


**Supplemental Figure 2.** Saline-paired context entries. (**A**) Heatmap of VTA-NAc activity for the 4 seconds surrounding the first 15 filtered entries into the saline-paired context. Color represents zΔF/F averaged across all VTA-NAc mice. (**B**) Same as A, but for VTA-PFC.
